# Spatiotemporal Evaluation of The Human Footprint in Colombia: Four Decades of Anthropic Impact in Highly Biodiverse Ecosystems

**DOI:** 10.1101/2020.05.15.098855

**Authors:** Camilo Andrés Correa Ayram, Andres Etter, Jhonatan Julián Díaz-Timoté, Susana Rodríguez Buriticá, Wilson Ramírez, Germán Corzo

## Abstract

The maintenance of biodiversity and the capacity of natural systems to provide goods and services for people is affected on different levels by the intensity of human activities on ecosystems. In this study, we apply a Legacy-adjusted Human Footprint Index (LHFI) to evaluate the spatiotemporal variation of anthropic impact in Colombia in 1970, 1990, 2000 and 2015. We identified hotspots of change in LHFI and we evaluated the intensity of anthropic pressures in natural regions and ecosystems. We found that LHFI in Colombia increased between 1970 and 2015. The Andean and Caribbean regions presented the highest levels of anthropic impact, remaining stable over time. Hotspots of change were mainly located in the following regions: Andean (Antioquia, Cauca and Valle del Cauca states), Amazon (Amazonas, parts of Meta, Guaviare and Putumayo states) and Orinoco (Casanare and parts of Meta and Vichada states). In addition, ecosystems that under the IUCN ecosystem risk categories are Critically Endangered (CR) and Vulnerable (VU) were the most affected by a high level of human impact. Spatiotemporal evaluation of the human footprint in Colombia provides new insights about trends in human pressures on ecosystems and constitutes an analytical tool with high potential for harmonizing land use planning and biodiversity conservation.

## 1. INTRODUCTION

Human pressures on the environment have drastically accelerated since the mid-twentieth century, risking biodiversity and the provision of goods and ecosystem services (Steffen et al., 2015). Direct impacts of human activities on natural systems include habitat loss and degradation (Crooks et al., 2011), fragmentation (Haddad et al., 2015), deforestation (Hansen et al., 2013), extinctions of species (Dirzo et al., 2014) and plastic pollution in marine ecosystems (Eriksen et al., 2014). Consequently, in the last two decades, landscapes that had remained almost free of human impacts suffered a reduction of one tenth of their surface, predominantly in highly biodiverse regions with pervasive socioeconomic inequality (e.g. the Amazon 30% and Central Africa 14%). This trend highlights the need for national and international prompt actions that recognize the conservation value of such areas and to manage them according to the unprecedented threats they face (Watson et al., 2016). To achieve this, it is key to understand spatiotemporal patterns of human impact on natural systems, an issue addressed by a number of studies (Etter et al., 2011; Sanderson et al., 2002; Venter et al., 2016; Woolmer et al., 2008).

One of the first indicators of human impacts on ecosystems was the Human Footprint Index (HFI, -Sanderson et al., 2002). HFI originally uses four spatial layers (population density, land transformation, accessibility and electric power infrastructure) but it has been modified to include more information (i.e. Woolmer et al., 2008 included mine sites and Leu et al., 2008 incorporated risk of exotic species invasion or anthropogenic fires). All HF indices are synthetic indicators that can be estimated at different scales depending on the homogeneity of the available information (Woolmer et al., 2008). In fact, HFI has been estimated globally to understand the human impact on the world’s biomes (Sanderson et al., 2002; Sanderson, 2013; Venter et al., 2016) and regionally (Tapia-Armijos et al., 2017; Trombulak et al., 2010; Woolmer et al., 2008) or nationally (in Colombia -Etter et al., 2011, in Mexico González-Abraham et al., 2015) to evaluate ecoregions or ecosystems.

Mapping spatiotemporal changes in human footprint reveals places where anthropogenic pressures have increased, decreased or remained stable, as well as, hotspots where impacts are outstanding (Geldmann et al., 2014; Li et al., 2018; Tapia-Armijos et al., 2017; Venter et al., 2016). Recently, HFI was used by Venter et al. (2016) to analyze global human impact changes between 1993 and 2009, while a number of recent studies show the practical use HFI to inform conservation planning (Correa Ayram et al., 2019, 2017; de Thoisy et al., 2010; Di Marco et al., 2013; Dobrovolski et al., 2013; Trombulak et al., 2010).

The most recent application of this approach for Colombia was proposed by Etter et al. (2011) whose HFI version includes the dimension of land use intensity, along with time of human intervention on ecosystems and their biophysical vulnerability (soil fertility, slope, moisture availability and number of short range species). By combining these three spatial dimensions, authors provided a more integral characterization of human impacts by incorporating historical and ecological contexts; this extended HFI facilitates ecosystem-specific detection of priority areas for conservation planning. In fact, it has already been used for this purpose in highly biodiverse countries, like Colombia (Ocampo-Peñuela and Pimm, 2014), China (Qiu et al., 2015) and México (Correa Ayram et al., 2019, 2017).

Despite its applicability, the extended HFI aggregates realized and potential human impacts and makes it difficult to compare human pressure in areas with different vulnerability factors. Incorporating vulnerability might undercover trends or patterns of human activities that need to be corrected. Conversely, by incorporating time since disturbance, the extended HFI version explicitly acknowledge that ecosystems might carry legacy effects from past landscape transformation that might not be evident before a tipping point is reached (Gardner et al. 2009).

In the last 50 years, ecosystem transformation in Colombia has been linked to the expansion of productive land and its technological changes in response to trade demands, increase in migration to urban centers, drug trafficking, and the internal armed conflict (Etter et al., 2008). In the last three years, the implementation of peace agreements between the government and the FARC-EP guerrillas has opened previously inaccessible areas, raising concerns about the expansion of deforestation and ecosystem degradation (Clerici et al., 2018; Negret et al., 2017).We offer a vision of human impacts that is as recent as possible, temporarily consistent, and it has a more precise spatial resolution (300 m) than the extended HFI calculated by Etter et al. (2011) or by any other global human footprint maps (e.g. Allan 2017, Venter et al., 2016). Thus, this study provides a trend baseline for monitoring human pressure on biodiversity and for designing prospective approaches (Trombulack et al., 2010).

In this study, we propose modifying the extended HFI (Etter et al., 2011) by only including time of human intervention. By including the cumulative impact of human actions, our Legacy-adjusted HFI (LHFI) provides a more conservative approach than current global HFI approaches (e.g. Venter et al. 2016, Sanderson et al. 2002), allowing the distinction of areas with recent but unprecedented interventions, from areas with a long history of human impact, where legacy effects might be stronger. The objective of this study three-fold: to conduct for the first time a multitemporal analysis of LHFI (1970-2015) for Colombia, to identify the hotspot of human footprint change, and to explore relationships of LHFI with ecosystem risk categories for every ecosystem. In contrast to other studies of human footprint (Li et al., 2018, Tapia-Armijos et al., 2017, Venter et al., 2016, Geldmann et al., 2014), we estimate human impacts in four periods which extends the temporal resolution of any available HFI.

## 2. METHODS

LHFI was calculated for continental Colombia (Eq.1), one of the most biodiverse countries in the world (Myers et al., 2000). Colombia comprises eight terrestrial biomes (Deserts and Xeric Shrublands, Mangroves, Paramo, Tropical and Subtropical Dry Forests, Tropical and Subtropical Grasslands and Savannas, Shrublands, Tropical and Subtropical Forests and Wetlands -Etter et al., 2015) and it is divided into six biogeographical regions (Andean, Pacific, Caribbean, Amazon, Orinoco and Catatumbo; Figure A-1).

In order to obtain a consistent LHFI estimations for time span of our study, we first followed the mapping procedure proposed by Etter et al. (2011): (a) Selection of the years of study and spatial resolution based on the availability of data and scale, (b) preparation of the six spatial variables for each year, (c) scaling the human pressure scores between 0 and 5, and (d) aggregate variables using equation 1, which results in four normalized (0-100) LHFI maps.

### 2.1. Calculation of the Legacy-adjusted Human Footprint Index

The LHFI was estimated using two of the three spatial dimensions proposed by Etter et al. (2011) as follows:

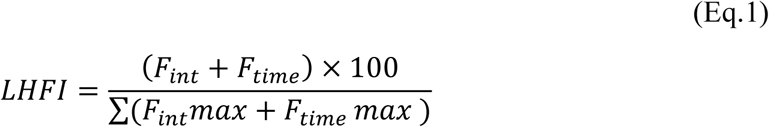

Where F_int_ is land use intensity and F_time_ is the time of human intervention on ecosystems. Etter et al. (2011) define F_int_ as the level of habitat modification due to extraction of resources and predominant land uses and management. F_time_ is the duration of time that the landscape has been subject to human activities, estimated from historical maps from the 1500’s, 1600’s, 1900’s and current maps from 1970 (see Etter et al., 2008 and 2011, Appendix 1). These two dimensions were estimated as follows:

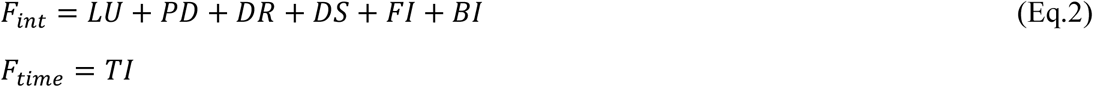

Where LU is land use type, PD is rural population density, DR is distance to roads, DS is distance to settlements, FI is the fragmentation index of natural vegetation, BI is the biomass index relative to natural potential, TI is time of intervention on ecosystems in years (see Etter et al., 2011 for a conceptual framework and Appendix 1 for preparation details of LHFI variables).

All seven primary variables were estimated at four years (1970, 1990, 2000 and 2015 -Table 1) and re-scaled between 0-5 to reflect their relative contribution to human impact and transformation; LGHI=0 indicates a null contribution and LGHI=5 indicates a very high contribution (Table 2). For each year, both F_int_ and F_time_ were estimated as the sum of its constituent variables and the final index was the normalized (between 0 and 100) to the sum of the two spatial dimensions (Eq.1). Value change analysis of LHFI was carried out for four periods: 1970-1990, 1990-2000, 2000-2015, and for the entire period from 1970 to 2015.

**Table 1.**
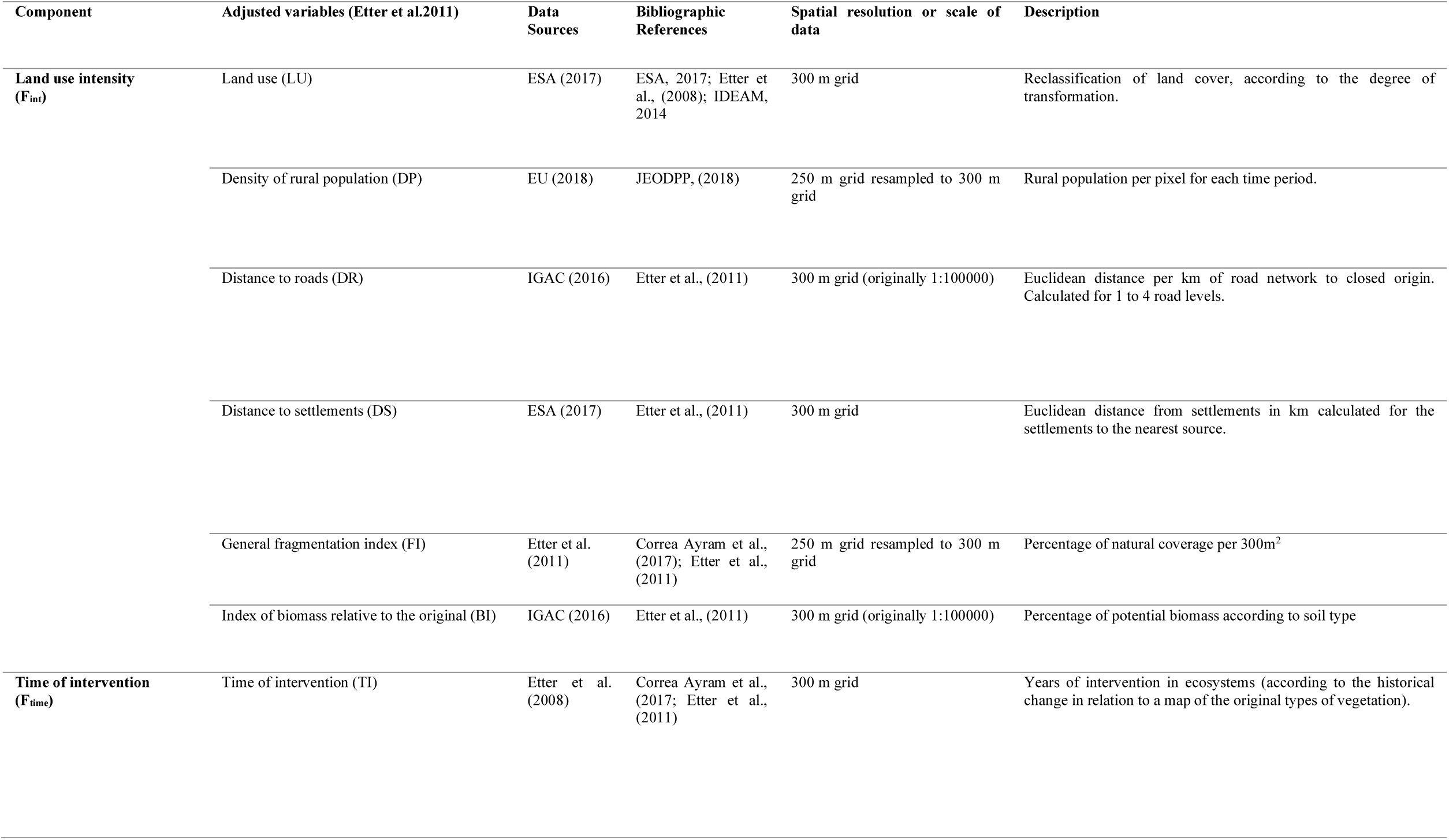
Data sources for construction of the two components of LHFI and their variables

**Table 2.**
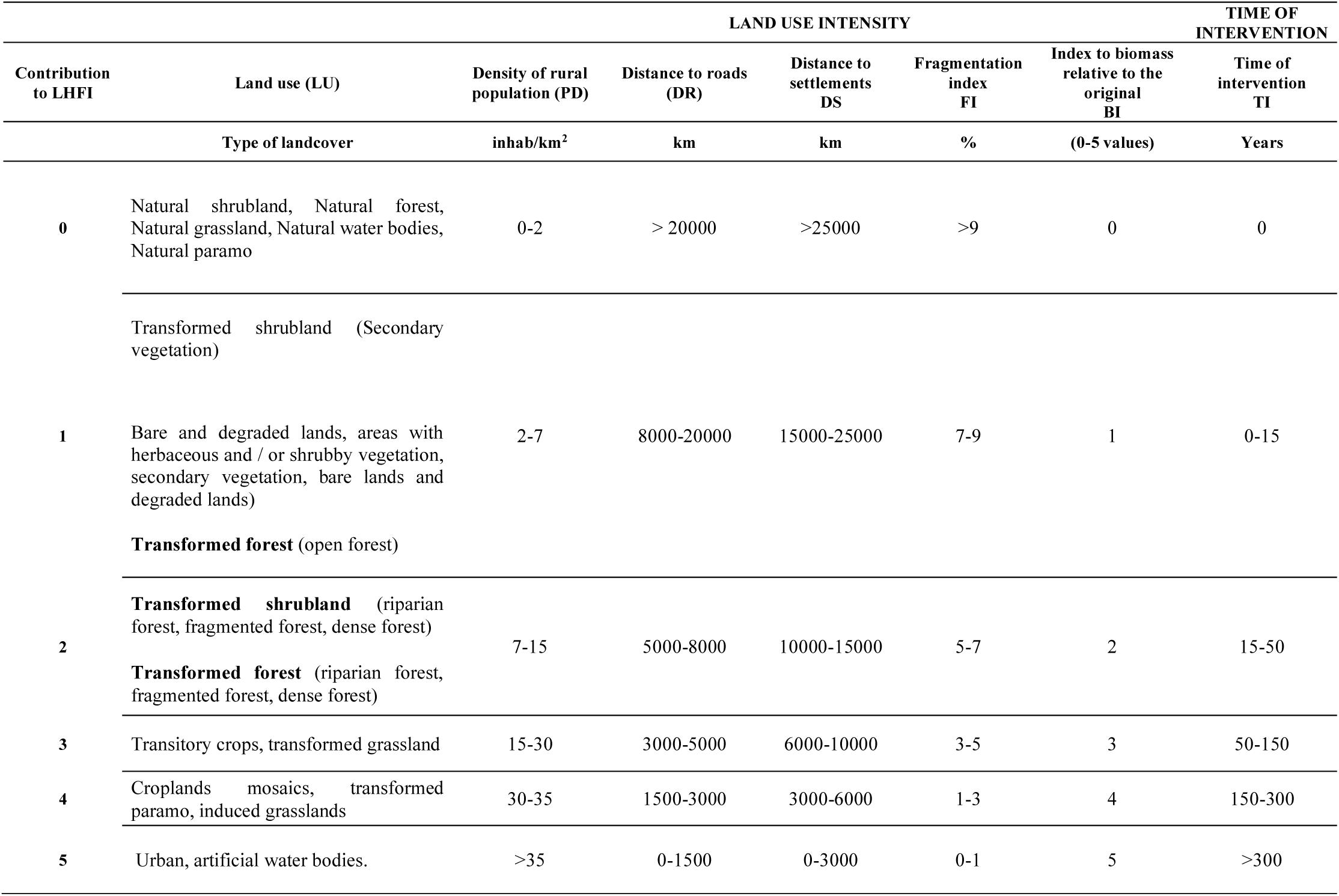
Contribution to LHFI and scaling of values ranges of each variable.

The study years were selected considering data availability and quality of geographic information; mainly binary maps of transformed -untransformed areas (Etter et al., 2008) and land cover maps (LC-CCI from ESA, 2017). Following Etter et al. (2008 and 2006), the period from 1970 to 1990 was characterized by strong population growth. From 1970, several important factors changed the Colombian economy and therefore the anthropic pressures on the biophysical landscape. First, there was the consolidation of the urbanization in major cities. Second, there was an increase in migration towards lowlands along the Andean-Amazon foothills. Third, the illegal economies around coca plantations showed an increase and a steady growth. Fourth, the armed conflict was increased due to the strengthened illegal economies. Fifth, parallel to these factors, a substantial change in policies toward the environment took place; these include the development of the Colombian Natural Resources Code, the growth of the National Parks System, and the recognition of indigenous and Afro-Colombian land rights occurred. Mainly, from 1990 to 2000 in cattle ranching had a dramatic expansion and some areas experience land abandonment (some underwent natural regeneration) due to the internal conflict (McAlpine et al. 2009). Discovery of new oil fields (e.g. “Cusiana” in Orinoco Region) promoted the increase in population density in historically sparsely populated areas. The expansion of livestock activity and oil infrastructure supported the rapid expansion of road constructions, particularly in the inter-Andean valleys (Palacio, 2001). From 2000 to 2015, cattle ranching continued to expand rapidly, growing in the lower areas of the Amazon region despite lack of improvements on beef prices and a decline in livestock incomes (Dávalos et al., 2014). A boom in mining and oil concessions characterized this period threatening fragile ecosystems such as floodplains and forests in Andean-Amazon foothills. The intensification of the armed conflict during 2010 and 2013 maintained relatively low levels of forest disturbance but this intensified during the negotiations period (2013-2017) and post-agreement period (2017-2019) (Murillo et al., 2020)

### 2.2 Change analysis and its relationship with ecosystem risk of collapse

To detect the areas with high or low LHFI change values, absolute change for each period was calculated as follows:

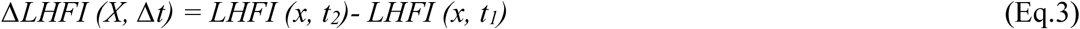

Where Δ*LHFI* (X, Δt) is the change in human footprint values during period X. *LHFI* (x, t2) is the human footprint values at time 2 and *LHFI* (x, t1) corresponds to the values at time 1. Δ*LHFI* categories were defined according to Tapia-Armijos et al. (2017) and Li et al. (2018) as follows: “Decrease” (Δ*LHFI* <0), “No apparent change” (Δ*LHFI* between 0 and 10), “Low growth” (Δ*LHFI* between 10 and 15), “Medium growth” (Δ*LHFI* between 15 and 30), and “High Growth” (Δ*LHFI* > 30). Values greater than 10 were classified as “hotspots of change” according to their intensity. We estimated the mean trend for each biogeographical region and for the entire country between 1970 and 2015 using simple linear regression models.

Similar to Tapia-Armijos et al. (2017), we used LHFI deciles to create the following human impact categories: “Natural” (LHFI of 0-15), “Low” (LHFI = 15-40), “Medium” (LHFI = 40-60) and “High” (LHFI> 60). Persistence of pixel classification between 1970 and 2015 were evaluated using the following categories: pixels that remained “Natural” were classified as “Persistence to low human impact” (PLHI); pixels that remained “High” were classified as “Persistence to high human impact” (PHHI); areas that remained “Intermediate” were classified as “Persistence to intermediate human impact” (PIHI).

We explored agreements between human impact categories and categories for IUCN ecosystem risk of collapse (Etter and Arevalo, 2016) by calculating a contingency table for each biogeographical region. The tables were used to estimate the percentage change in area between periods and for each combination of IUCN ecosystem risk and LHFI-based human categories.

All data were adjusted to raster data of 300 m resolution using MAGNA-SIRGAS / Colombia Bogota zone EPSG projection; LHFI calculations and spatial analyses were conducted in ESRI ArcGIS 10.7 and statistical analysis were conducted with R version 3.5.3.

## 3. RESULTS

### 3.1 Spatiotemporal variation of human footprint

In Colombia, the Legacy-adjusted Human Footprint Index (LHFI) increased 50% between 1970 and 2015 (Figure 1) and natural areas reduced to less than half of the national territory (for more details see Table A-3 in Appendix 1), although variation is high among regions. The Caribbean and Andean regions, which have the highest population densities showed the highest degree of LHFI, while larger regions with lower population densities (Amazonia, Orinoco and Pacific, with densities of 5 to 17 people / km2 -Etter et al. 2011), have more natural areas and lower LHFI values (Figure 1). In the Caribbean region, plains and alluvial valleys along major rivers (Magdalena, Sinú, San Jorge) showed the highest LHFI and the smallest proportions of natural areas. In the Andean region, which has the highest proportion of areas with high LHFI values (high LHFI values changed from 3% to 5.6% between 1970 and 2015), are concentrated in the inter-Andean valleys and along the Eastern Cordillera foothills (Figure 2).

**Figure 1.**
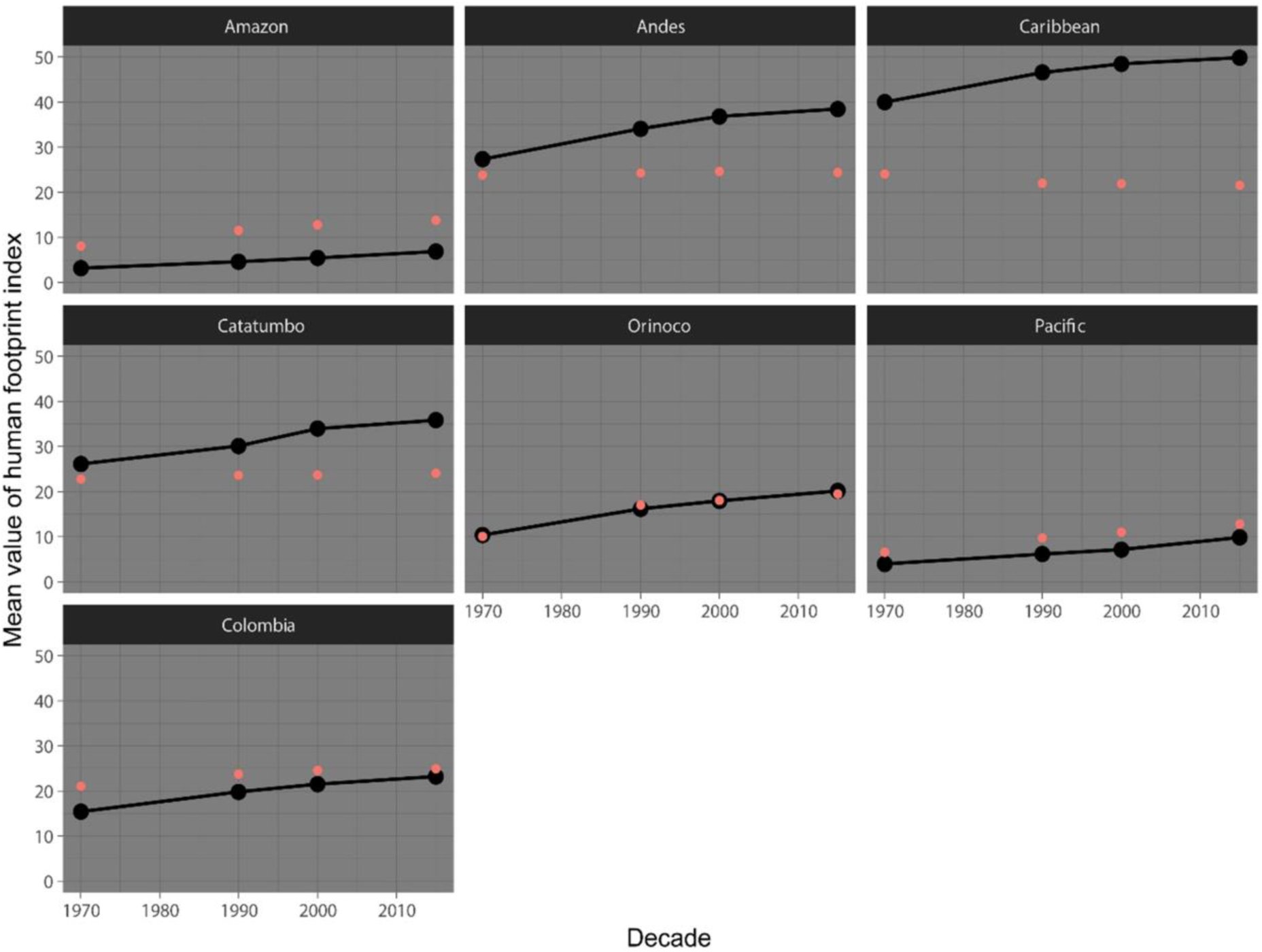
Distribution of mean values of LHFI in Colombia and its natural regions in 1970, 1990, 2000 and 2015. Red dots show standard deviation from the mean.

**Figure 2.**
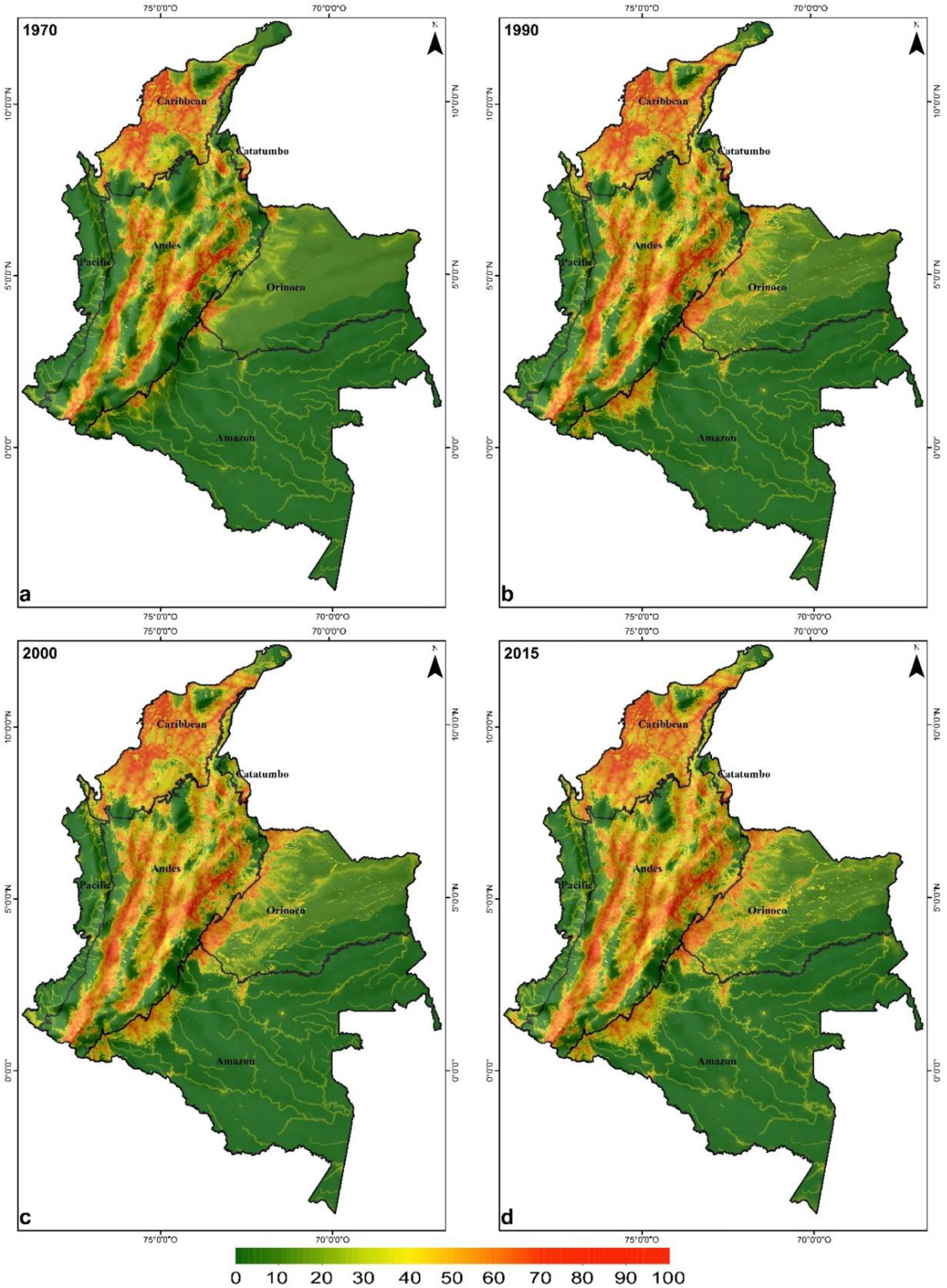
The maps show the spatial distribution of human footprint values in Colombia in each year of study. Spatial resolution 300m X 300m

The temporal dynamics of LHFI shows that, half of the country remains under low human impact and has persisted as such for at least 45 years (54.5% under PLHI category, Figure 3). Most PLHI areas are in the south and southeastern of Amazon region, southeastern of Orinoco region, and northwestern of Pacific region. A lower percentage of the country (38.2%) showed persistence of intermediate human impact (category PIHI) and most areas are north of the Andean region, south of the Caribbean region, and at the center of the Orinoco region. Some of these areas are currently associated with deforestation fronts (e.g. along the Eastern Cordillera foothills: Andean-Orinoco and Andean-Amazon foothills). Areas that remained under the high impact category (PHHI) correspond to 7.2% and concentrate in areas with agricultural and livestock production, such as the inter-Andean valleys, the Andean-Orinoco foothills, and the Caribbean coast.

**Figure 3.**
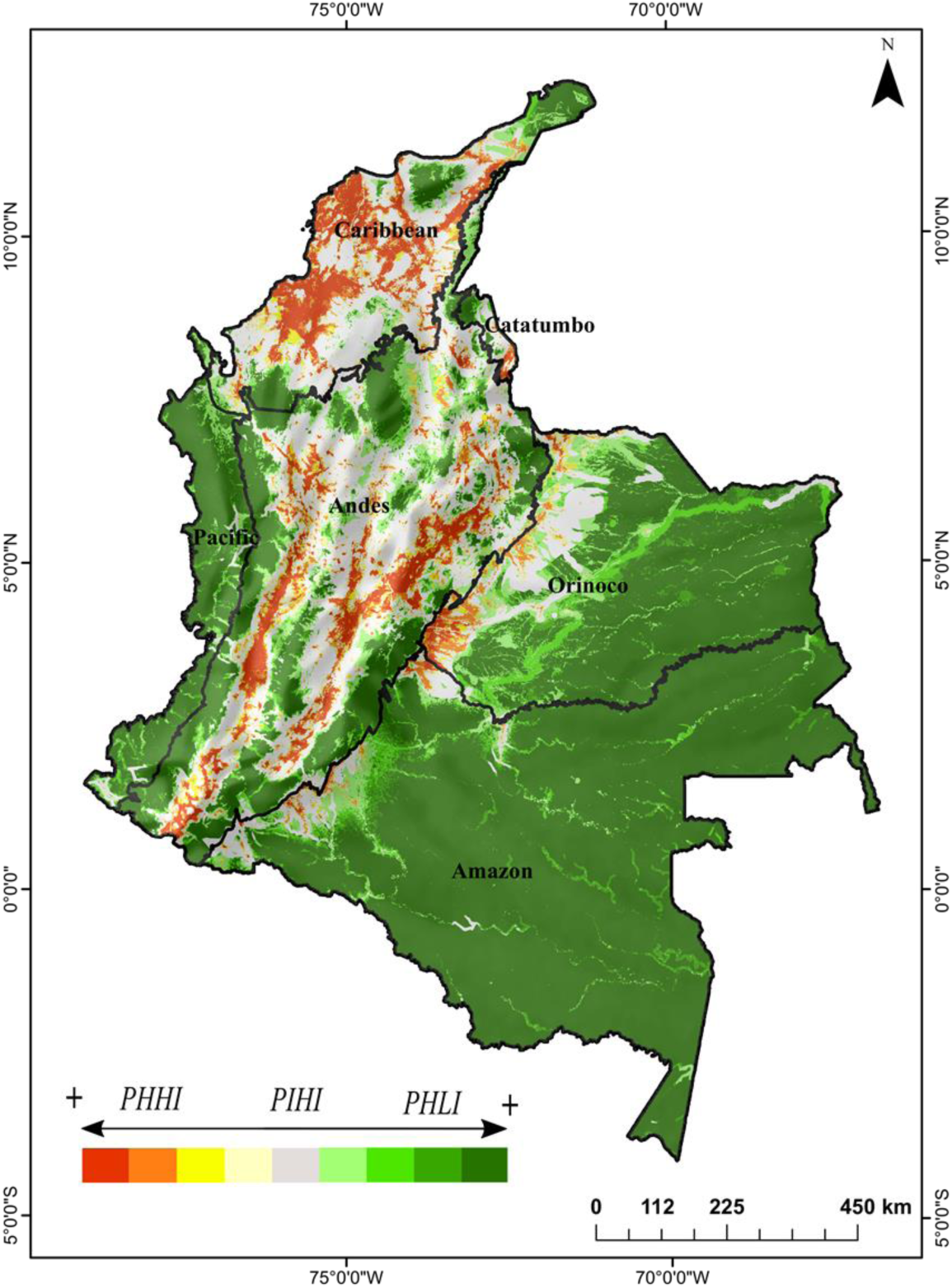
Gradient of high and low LHFI persistence between 1970 and 2015. Warm colors indicate areas with high persistence of high LHFI (PHHI) and green colors persistence of low LHFI values (PLHI). Gray indicates areas that were at intermediate values during the study period (PIHI).

At the national scale, the highest LHFI change occurred between 1970-1990 (28.3%) while spatially, the Orinoco and Pacific regions had the most notable change during the entire study period. Between 1970 and 1990 the greatest change concentrated in the Eastern Cordillera foothills (Orinoco region), Darién subregion (in the northwestern Pacific region), central valleys of the Andean region, northeastern Caribbean region, and Serranía de San Lucas (at the northern end of the Central Cordillera -Figure 2a and Figure 2b). For the entire study period, prominent changes were evident along the Andean-Orinoco foothills and towards the southwest in the Pacific region where annual growth rates were 2% (Figure 2d and Table A-5 in Appendix 1).

Hotspots of change in LHFI concentrate in the Andean, Caribbean and Orinoco regions (Figure 4). The period of 1970-1990 reflects the most dramatic changes, in all regions except the Pacific and Catatumbo (Figure 4a), and this intensity remains once we considered the different period duration (see annual increment estimations in Appendix 1-Table A-5). The 1990-2000 period shows more stability compared to the other periods; yet, new hotspots did emerge in the southwestern Andean, southeastern Orinoco, southwestern Amazon, and the central Catatumbo regions (Figure 4b).

**Figure 4.**
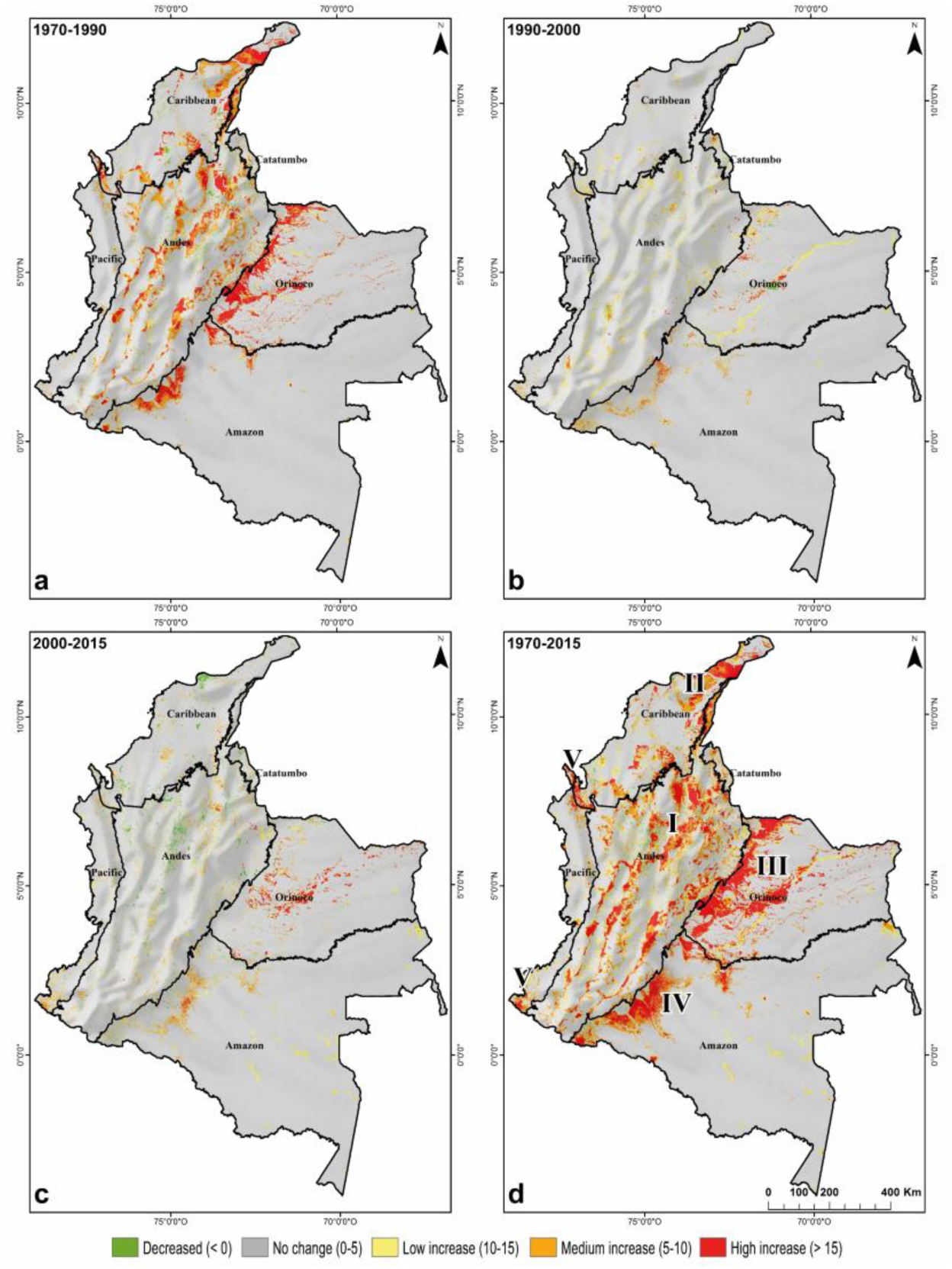
Spatial distribution of hotspots of change for each year evaluated. a) Period 1970-1990. b) Period 1990-2000. c) Period 2000-2015. d) Period 1970-2015. Between 1970 and 2015, five main change hotspots were identified: I) North-central Andean region. II) Lower part of the Sierra Nevada de Santa Marta and the peninsula of La Guajira. III) Andean-Orinoco foothills. IV) Andean-Amazon foothills at the deforestation front of Caquetá and Putumayo States. V) Darién sub-region in the Chocó State. V) Southwestern Pacific region.

The 2000-2015 period (Figure 4c) shows an area with concentration of high LHFI values consolidated in the Amazon region corresponding to the deforestation front of the states of Caquetá and Putumayo, which expands to the Orinoco region. Hotspots located along the Andean-Orinoco foothills, show an increase in concentration of high LHFI values in the southeastern Orinoco region. In general, 73% of the Orinoco region remained with low LHFI levels during the studied period; however, a decrease of the area (7%) with low LHFI values (15-40) between 1970 and 2015 was detected. New hotspots also appear in the southwestern and central Pacific region. Areas of LHFI decrease were few and more prominent between 2000 to 2015 (Figure 4c). These areas are located mainly to the north- and southwest parts of the Andean region, and the southern Caribbean region (areas bordering the Sierra Nevada de Santa Marta).

### 3.2 Human impact and threatened ecosystems

Between 1970 and 2015, all IUCN ecosystem risk categories had an increment of areas with medium or high LHFI values (Table 3). The category of Critical Endangered (CR) had the highest proportion of medium and high LHFI areas; especially for ecosystems such as dry tropical forest, tropical desert, the intrazonal dry ecosystems of the Andean region, humid tropical forests of the Andean-Orinoco foothills (Orinoco region), and wetlands in the central-eastern Andean region (states of Boyacá and Cundinamarca). Within the Endangered category (EN), natural savannas within the eastern plains (Orinoco region) increased its percentage of high LHFI values. For Vulnerable ecosystems (VU) there was considerable increase of medium to high LHFI areas until 2000, and then a decrease; especially for the Paramo ecosystem. Least Concern (LC) ecosystems such as humid tropical forest of Amazon region, had the largest representation of low LHFI values. However, this ecosystem within the Amazon showed a rapid increase in LHFI values, especially in areas clustered along the Andean-Amazon foothills, which coincides with deforestation hotspots.

**Table 3.** LHFI percentage by category of ecosystem threat. In bold, the highest percentages for each category of threat are highlighted, according to the LHFI evaluated.

| IUCN Red List categories | LHFI | 1970 | 1990 | 2000 | 2015 |
| --- | --- | --- | --- | --- | --- |
| CR | Natural | 2.41% | 1.23% | 1.06% | 0.98% |
|  | Low | 3.44% | **6.44%** | 1.17% | 1.02% |
|  | Medium | **4.62%** | <u>1.54%</u> | <u>5.97%</u> | <u>5.80%</u> |
|  | High | <u>4.50%</u> | <u>5.76%</u> | **6.76%** | **7.18%** |
| EN | Natural | **3.72%** | <u>2.35%</u> | 2.08% | 1.87% |
|  | Low | 2.17% | 1.42% | **2.53%** | **2.37%** |
|  | Medium | 0.70% | **2.58%** | 1.59% | 1.78% |
|  | High | 0.28% | 0.52% | 0.67% | 0.85% |
| VU | Natural | **13.76%** | **11.39%** | **10.62%** | **10.06%** |
|  | Low | 4.22% | 5.31% | 4.30% | 4.31% |
|  | Medium | 3.37% | 4.34% | 5.59% | 5.84% |
|  | High | 0.92% | 1.23% | 1.76% | 2.06% |
| LC | Natural | **50.35%** | **47.33%** | **45.00%** | **42.19%** |
|  | Low | 3.89% | 2.98% | 6.52% | 8.37% |
|  | Medium | 1.31% | 4.91% | 3.44% | 4.17% |
|  | High | 0.33% | 0.66% | 0.93% | 1.15% |

## 4. DISCUSSION

Mapping LHFI at four different periods improved our understanding of how land transformation acting at different spatiotemporal scales translate into a footprint with ecological consequences. First, our results show that regions that have historically promoted productive activities and that have been at the center of Colombia’s development such as the Andean and Caribbean regions, have persistently high LHFI locations (PHHI). These regions also contain highly dynamic areas and this mixture redound in large spatial and temporal variability of LHFI values. Both regions represent regions with prevalence of agriculture that since the 1970’s have experienced continuous population growth and urban industrialization (Etter et al., 2008). Second, areas with a more recent history of transformation are already experiencing localized and persistent high human impact, while showing variable impacts within a matrix of persistently low LHFI values; this is the case of the Orinoco and to a lesser degree, the Amazon region (Figure 2). In these regions, changing areas in terms of LHFI are in the eastern plains and along the Andean foothill (Figure 2), which are areas of active agricultural expansion. Between 1970 and 1990 in these areas, cattle ranching expanded along with the use of exotic forages (Etter et al., 2008) and since the end of the 1980s, African oil palm plantations has also expanded (e.g. 33 km^2^ in 1987 to 162 km^2^ in 2007). In the Orinoco region, LHFI increment might also be due to changes in population density that had increase since 1970 and had a peak increment during the petroleum boom (1985-1993; 0.01 inhab/km^2^ in 1970 to 70 inhab/km^2^ in 2000-Romero -Ruiz et al. 2012). All these changes go hand and hand with an intense road network expansion between 1970 and 2000.

In contrast to areas with gradual increase of LHFI, specific areas in regions with low population density, have recently and rapidly increased their LHFI values (between 2000-2015). These areas are the Andean-Amazon foothills in the states of Caquetá and Putumayo, northwestern Amazon lowlands in the state of Guaviare, and certain zones within the Pacific region. The first two are current deforestation hotspots (Armenteras et al., 2013; Dávalos et al., 2014; Etter et al., 2006; Hoffmann et al., 2018) that experienced colonization processes since 1960 but have recently had high immigration rates (Dávalos et al., 2014). Andean-Amazon foothills have also been impacted by low-scale mining, often illegal, since the time their colonization began (Hoffmann et al., 2018). In the Pacific, land cover changes can be explained by mining for gold and precious metals (silver and platinum; Servicio Geológico Colombiano, 2012) alongside with expansion of African oil palm plantations. Illegal coca expansion is another contributing factor to fast changing LHFI. Since the 1980’s (Armenteras et al., 2013; Dávalos et al., 2014), coca crops have increased in the Andean region, but they are more recently affecting the Pacific region (Dávalos et al., 2014; Rincón Ruiz et al., 2013). According to Rincón Ruiz et al. (2013), the expansion occurred between 2001 and 2008 when coca crops went from representing 1% to 11% of the region.

Should the increase pattern in LHFI continue, consequences for biodiversity could be devastating. Many of the most biodiverse ecosystems in the planet are in Colombia, which holds 10% of global biodiversity (Myers et al., 2000). Since 1970, the human footprint has increased 50%, an alarming situation considering that 65% of the Colombian ecosystems are now threatened (Etter and Arevalo, 2016) and that the global average increment for tropical biodiverse areas is estimated at 20% (Venter et al., 2016). Like ours, other studies highlighted that most endangered ecosystems are located in human dominated landscapes (Brauneder et al., 2018; Cincotta et al., 2000; Venter et al., 2016), which is not surprising because IUCN criteria, include variables related to human pressure (Keith et al., 2013, Etter and Arevalo, 2016). Beyond this expected pattern, the spatial agreement implies that urgent actions are mostly needed not in almost intact areas, but in regions with high human impact and good representation of unique ecosystems. The Andean region holds a large number of endemism (Ocampo-Peñuela and Pimm, 2014), it has been prioritized for biodiversity conservation (Myers et al., 2000; Orme et al., 2005; Anderson and Maldonado-Ocampo, 2011), and holds the Colombian paramos, a key ecosystem for hydric regulation and water supply (Cárdenas et al., 2017; Etter et al., 2011). Similarly, increase human impacts along the Caribbean region threatens wetlands and the endangered dry tropical forest, which have been historically transformed to favor agricultural expansion (Aldana-Domínguez et al., 2017; Márquez and Colaboración de Pérez García, 2001; Patino and Estupinan-Suarez, 2016). Although we detected that human footprint decrease in some dry tropical forest areas in the Caribbean region, this can be caused by changes in the dynamics of rural settlements (Schubert et al., 2018). Further research is required to evaluate drivers of land cover change, especially because this ecosystem has the most cumulative impact of human activities during our study period and historically, it is one of the most impacted (Ferrer-Paris et al. 2019).

Increments in LHFI values in other areas such as the Andean-Orinoco foothills and the Pacific region are also alarming because both regions are widely recognized by their impressive and unique biodiversity (Myers et al., 2000; WWF, 2014). Although the tropical rain forest is not endangered, deforestation expansion along the foothills and Amazon lowlands is fragmenting a habitat which could interrupt functional connectivity between the Andes and the Amazon (Clerici et al. 2018). Likewise, current transformation in the Pacific Region endangers an area considered to be a global biodiversity hotspot that functions as a biological corridor between the southeastern portion of Panamá in the Chocó -Darién subregion and the pacific coast of Ecuador and Peru. Our results provide an opportunity to evaluate different land management practices, for example, between indigenous reserves and collective territories of Afro-Colombian communities with areas outside these territories. These territories cover 50% of the Pacific region, representing a fortress for biodiversity that along protected lands could contain human pressures in the future (Dávalos et al. 2011, Cámara-Leret et al., 2016).

A major limitation of the study is its spatial resolution that might limit its use for official purposes. Given that the main objective was to conduct a multitemporal analyses, we prioritized data that could be compared during the study period. Official land cover datasets come with a higher spatial resolution (∼30 m) but they are only available after 2000 and prior to 2013. To facilitate the use of our procedures for official purposes, we harmonized our land cover classification using the official legend (IDEAM, 2010). Information limitations also apply to primary socioeconomic and human activity variables in areas with low land use intensity. It was not possible to have good quality data on economic activities and poverty indices, communication antennas, garbage dumps, mining, soil for agriculture, fire occurrences, hunting data, among others (Geldmann et al., 2014). Lacking this information limits the ability of our methods to detect human impacts in the Amazon, Pacific, and Orinoco regions, despite of the general knowledge about the impacts that small-scale mining is causing there. Therefore, our results might underestimate recent changes in these regions. Finally, we could only calculate cumulative changes for periods lasting 10, 15, and 20 years; which limits our ability to fully characterize the underlying processes producing high LHFI, their interactions, and their local scale signatures.

Despite these limitations, one of the most relevant applications of our work is the monitoring of LHFI in the context of Colombia’s armed conflict and Peace Agreements effects (Colombian Government and FARC -EP 2016). Since the signing of the Peace Agreements, post-conflict dynamics have favored the expansion of human activities by reversing of long standing, low impact land management enforced by FARC-EP guerrillas; many territories are suffering the negative effects of recolonization (Clerici et al., 2018; Negret et al., 2017). A recently study by Murillo et al. (2020) found that during the post-peace agreement period (2017-2018), in the LHFI hotspot of Andean-Amazon foothills (see hotspot IV in Figure 4) forest disturbance increased by 50% over the estimate of 2013-2016 (Hoffmann et al., 2018). In general, deforestation associated with the return of rural population (Álvarez, 2003; Baptiste et al., 2017) is increasing human pressure in areas with low historical values, such as Serranía de San Lucas, Catatumbo region or Serranía de la Macarena (Reardon, 2018). In a scenario of successful implementation of the peace agreements, greater LHFI can be expected in rural areas due to the return of displaced populations and the reactivation of local economies (Negret et al., 2017). In contrast, rural areas with lower LHFI values that have reduced their internal armed conflict, may have potential for sustainable economic development (e.g. avitourism), as suggested by Scott and Ocampo Peñuela (2018).

Finally, LHFI can readily be used in landscape connectivity studies (Correa Ayram et al., 2017) to identify management priorities that integrate elements of the landscape (e.g. corridors, restored patches, among others) and encourage connectivity of multiple species (Correa Ayram et al., 2019). This approach is key for public policies that regulate land use in priority areas for ecosystem connectivity, as it is for deciding where to establish such areas to effectively achieve global conservation goals in the first place (e.g. Aichi Target 11) (Saura et al., 2017). The multitemporal character of our work provides the bases for prospective models that identify areas with a higher probability of human footprint increase. The challenge would be to incorporate climate change scenarios to propose adaptive strategies that include aspects of the impact of land use and land cover change at local and regional scales (Nuñez et al., 2008; Salazar et al., 2016; Swann et al., 2015).

## 5. CONCLUSIONS

This study is the first spatiotemporal evaluation of human pressure in Colombia and it provides the first historic examination of where and how different levels of human impact are distributed. By considering time of intervention (TI), our Legacy-adjusted Human Footprint Index (LHFI) allows evaluations over realized human impacts without considering biophysical vulnerability as previously suggested by Etter et al. (2011). Without this dimension, our approach distinguishes areas that show high human impact due to recent degradation process from areas with historical human pressure. This attribute is essential to identify areas where human pressure is so prevalent that ecosystems could be beyond the tipping point for recovery (Pimm et al., 2006; Ceballos et al., 2015), or areas where natural recovery and induced restoration processes are more likely to succeed (Jones et al., 2018).

Our study provides an evaluation of human impacts in a mega-diverse country and it allowed us to know where impacts have increased, decreased, or being maintained in the last 45 years. These estimations suggest three major regional and subregional patterns: the first pattern corresponds with regions of long history of human impact at the center of the country’s development but where critically endangered ecosystems are at their highest vulnerability. A second pattern correspond to development front concentrated in the Orinoco region with intense transformation throughout highly biodiverse foothills due to expansion of low productive cattle ranching and an intensified agroindustry. The last pattern corresponds to areas of recent fast and intense land transformation associated with deforestation fronts and reactivation of local legal and illegal activities

We believe our approach, through a periodic update of LHFI has potential to evaluate restoration, invasive species control, monitoring conservation strategies, and the role of protected areas to contain human pressures. It can also be applied to other scales, landscapes or countries, substantially contributing to future evaluations of human impacts for better decision-making in biodiversity conservation planning.

## ACKNOWLEDGMENTS

The authors thank the Ministry of the Environment of Colombia for providing financial support to this research (Resolution 0130 of 2018). We recognize the valuable contributions of reviewers who helped improve the manuscript. We also want to thank Julia Premauer for their support in the revision of English version of the manuscript.

