## Appendix 1 for "Spatiotemporal Evaluation of The Human Footprint in Colombia: Four Decades of Anthropic Impact in Highly Biodiverse Ecosystems"

The following information provides detailed methods and synthetic results from primary variables used in the LHFI estimations.

1. **Study area**

We used continental Colombia as a case study to compare the changes in the accumulated human impact on ecosystems. Colombia (1.1 million km^2^) is in the northwestern part of South America, latitude (4°S and 12°N) and longitude (67° and 79°W). It has a large elevation range (0-5800 m) and variation in mean annual precipitation (from 150 mm in La Guajira peninsula to 12.000 mm in parts of the Chocó state). The mean annual temperature decreases with higher altitudes forming six distinct temperature zones: warm (0-1000 m; > 24° C) covering 91% of the country, temperate (1000-2000 m, 18-24°C) covering 5%, cold (2000-3000 m, 12-18°C) covering 2%. The remaining 2% (3000-5800m) has very cold weather (6-12°C) and comprises: sub-paramo (3-6°C), paramo (1.5-3 °C) and nival zones (<1.5 °C -IDEAM, 2005).

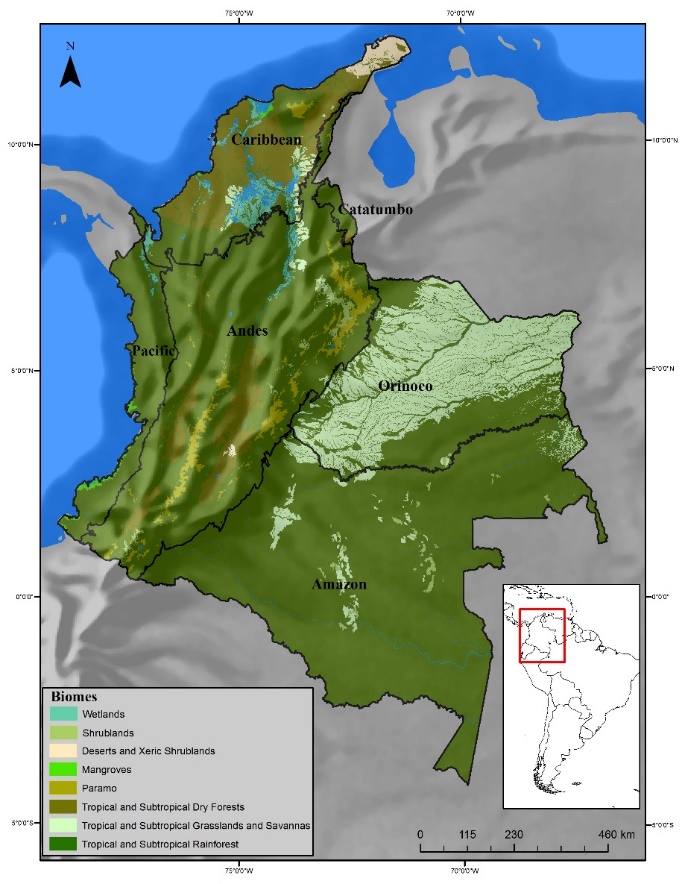

Figure A-1. Study area location showing the spatial distribution of Colombia’s main terrestrial biomes (Etter et al., 2015).

2. **Preparation of LHFI variables:**
1. **Spatial dimension: Land use intensity (F*_int_*)**
   a. **Land use (LU)**

Human land use change is essential to determine habitat loss and the related impact that differs among land uses (Etter et al., 2011; Woolmer et al., 2008). To obtain land use maps for 1970, 1990, 2000 and 2015, three datasets were used: 1) map of Colombian Potential Ecosystems (Etter et al., 2017), 2) Binary Maps of Colombia’s transformed-untransformed areas (T/UT) for the same years (Etter et al., 2017) and 3) Land cover maps of European Space Agency (LC-CCI from ESA, 2017) for the years 1992, 2000 and 2015. We harmonized land cover categories following Corine Land Cover methodology for Colombia (IDEAM, 2010) and the land cover legend in ESA LC-CCI products, which are based on FAO (2002). The three datasets were combined to get the transformation level for each ecosystem in each year of study, as well as the existing land covers.

To obtain the land cover map for 1970 we used 1970’s binary map (T/UT; Etter et al., 2017), the map of potential ecosystems (Etter et al., 2017) and the ESA land cover map for 1992. Natural areas on the 1992’s ESA map that were untransformed in 1970 were assigned to the equivalent potential ecosystem. Similarly, 1990’s cover map was estimated from 1990’s T/UT maps and ESA land cover map from 1992. For 2015 we used the ESA land cover categories for that year. Finally, we obtained nine categories for each of the years evaluated (1970-1990-2000-2015 -Table 2 in the main text for categories). Each land use category obtained was assigned a human footprint value from 0 to 5, where 0 is the least human footprint impact and 5 the highest impact, similar to Tapia-Armijos et al. (2017).

b. **Distance to roads (DR)**

Roads affect the landscape ecological processes through habitat loss and fragmentation, increase of edge effect, reduce landscape permeability and reduce habitat quality (Carr et al. 2002). Roads also cause direct impacts on wildlife such as roadkill for vehicle collision, which has been the most widely recognized impact of roads on biodiversity in the last three decades (Bennet, 2017).

Distance to road maps were obtained from roads distribution dataset available for the years 1970-1990-2000-2015. In the absence of an official dataset at national level before 2000, we reconstructed the roads maps for 1970 and 1990 from digitalization process of the aeronautical navigation charts available in “Perry-Castañeda Library, Map Collection of Colombia” of The University of Texas at Austin (2015). In order to guarantee a consistent and standardized dataset of roads from year to year, we used the most recent map (2015) as the starting point and eliminated roads that did not coincide with the dataset of aeronautical navigation chart of the same year. Road maps for most recent years (2000 and 2015) were built, based on official maps of Instituto Geográfico Agustín Codazzi (2016). All 4 maps only included road types 1, 2, 3 and 4 that correspond to roads that allow vehicle circulation (Table A-1). Finally, we used a simple Euclidean measure to map distance to all roads of each year of study. We assigned a human footprint scores, using the scaling values of Etter et al. (2011) from 0 to 5, indicating a lower to higher contribution to the degree of human footprint (for more details of HF range values see Table 2).

| **CATEGORY** | **TYPE** | **ROAD CHARACTERISTICS** | **SPECIFICATIONS** |
| --- | --- | --- | --- |
| 2 (two) or more lanes | 1 | Hard lining, Paved road | Greater than 5.5 meters wide |
| 2 (two) or more lanes | 2 | Loose or light coating, unpaved road | Greater than 5.5 meters wide |
| 1 (one) lane | 3 | Hard lining, Paved road | Between 2.5 meters wide and less than  5.5 meters |
| 1 (one) lane | 4 | Loose or light coating, unpaved road | Between 2,5 meters wide and less than 5,5 meters |

**Table A-1.** Road types categorization. IGAC (2016).

c. **Distance to settlements (DS)**

To map the influence of human settlements, we extracted the "urban areas" category of the land use maps of each year of study (ESA, 2017). This class describes areas that have an artificial cover as a result of human activities such as construction (e.g. cities, towns, and paved lands). We calculated the Euclidean distance between each human settlement. The final maps of distance to settlements (300 m resolution) for 1970, 1990, 2000 and 2015, were grouped following the same scaling as Etter et al. (2011) and classified in values from 0 to 5, indicating a lower to higher contribution to the degree of human footprint based on the classes shown in Table 2.

d. **Population Density (PD)**

To map the influence of population density, we used the data series of 1970-1990-2000 and 2015 developed by the JRC Earth Observation Data and Processing Platform (JEODPP). The data series summary the spatial distribution of population density at 250 m x 250 m gridded. The population density maps were resampled to match the 300 m resolution of the other data sets. The values of population density (Number of people / per pixel) were grouped following the same scaling as Etter et al. (2011) and classified in values from 0 to 5, indicating a lower to higher contribution to the degree of human footprint based on the classes shown in Table 2.

e. **Fragmentation Index (FI)**

To map the contribution of habitat fragmentation in terms of the human impact, we used the Binary Maps of Colombia’s transformed - untransformed areas for the years 1970, 1990, 2000 and 2015 (Etter, 2017) at 250 m resolution. The binary maps were resampled to match the 300 m resolution of the other data sets. To obtain continuous information that reflects the distribution of habitat fragmentation throughout the territory, we calculated the proportion of natural vegetation per pixel (300m) using a Kernel density method. The high-density values of natural vegetation per pixel show a low contribution to the human footprint. In contrast, low density values correspond to areas with high fragmentation and therefore a high contribution to the human footprint (for more details see Table 2).

**f. Biomass index (BI)**

An indicator of land use type and intensity includes the reduction of biomass through land cover transformation (Erb et al. 2018; Etter et al. 2011). We assigned a score according to the proportion difference in biomass between a natural ecosystem that should be in an area and the cover at that area due to transformation. For example, an original forest transformed into pastures presents a drastic change in the biomass contribution with respect to the original, therefore, a high contribution value is assigned to LHFI. In contrast, if the original forest is currently maintained, its biomass contribution remains high, which means a low anthropic impact and a low value of the human footprint. This procedure has already been applied by Correa et al. (2017) and it is adjusted based on Etter et al. (2011) for this study as follows: we obtained the biomass index map using the adapted land use maps for the years 1970-1990-2000 and 2015 at 300 m resolution (section a) and the potential ecosystems map of Etter et al. (2017). Based on the experts' criteria, a decision matrix was generated for the allocation of the human footprint values according to the availability of biomass in each ecosystem and classified in values from 0 to 5. For this task we considered the 9 categories of land use obtained (See Table 2 for land use categories and BI contribution to LHFI) before as well as the attribute of potential ecosystems. Table A-2 shows the final matrix obtained with the human footprint values for each ecosystem evaluated.

2. **Spatial dimension: Time of intervention (F_int_)**

The time since humans have used the territory and the intensity of intervention affect the resilience of natural systems and are factors that determine the magnitude of the accumulated anthropic effect on the landscape (Correa et al., 2016). TI corresponds to the duration in years that landscapes and ecosystems have been subjected to human disturbance (Etter et al., 2011). To map the time of intervention (TI), we followed the method presented by Etter et al. (2011) based on assigning of years of human intervention from historic land use maps of Colombia constructed in the 1500’s, 1600’s and 1900’s. The years of intervention for 1970, 1990, 2000 and 2015 were estimated using the binary maps (T/UT) generated by Etter et al. (2017) for the same periods of time. The binary maps were combined to identify areas that have been subject to historical use. Subsequently, the years of intervention are added to obtain TI for each area. For example, all transformed areas for 1970 that were untransformed in 1920, were assigned 50 years of TI. Areas that remain untransformed since 1500 have the minimum TI of 0 and areas that were labelled as transformed in 1500 had at least 515 years of transformation. TI was classified in 6 (0 to 5) categories based on Etter et al. (2011-Table 2), assuming that a longer TI can generate a greater human impact on the landscape by interacting with the land use intensity (LU).

Table A-2. Matrix of allocation value of human footprint according to ecosystem and category of use for biomass index

| **Land cover types** | **Natural shrubland** | **Transformed shrubland** | **Natural forest** | **Transformed forest** | **Artificial water bodies** | **Natural water bodies** | **Transitory Croplands** | **Natural grassland** | **Transformed grassland** | **Mosaic of croplands** | **Induced grasslands** | **Urban** |
| --- | --- | --- | --- | --- | --- | --- | --- | --- | --- | --- | --- | --- |
| **Potential Ecosystem** |  |  |  |  |  |  |  |  |  |  |  |  |
| **Open and succulent shrubs of the Zonobiome of Tropical Deserts** | 0 | 2 | - | - | 5 | 0 | 3 | - | - | 3 | 4 | 5 |
| **Low and open shrublands and desert areas of the Zonobiome of Tropical Deserts** | 0 | 2 | - | - | 5 | 0 | 3 | - | - | 3 | 4 | 5 |
| **Xerophytic shrublands of the Zonal Orobiomes of the Zonobiome of Tropical Rainforest** | 0 | 2 | - | - | 5 | 0 | 3 | - | - | 3 | 4 | 5 |
| **Shrublands and Low dense Forests Sclerophyllous of the Pedobiomes of the Zonobiome of Tropical Rainforests** | 0 | 2 | - | - | 5 | 0 | 3 | - | - | 3 | 4 | 5 |
| **Shrubland and dense cardonal of the Zonal Orobiomes of the Zonobiome of Tropical Rain Forest** | 0 | 2 | - | - | 5 | 0 | 3 | - | - | 3 | 4 | 5 |
| **High dense mangroves of the Helobiomes of Zonobiomes of Tropical Rainforest** | - | - | 0 | 3 | 5 | 0 | 4 | - | - | 4 | 4 | 5 |
| **High and Dense Forests of the Holobiomes of the Tropical Rainforest Zonobiome** | - | - | 0 | 3 | 5 | 0 | 4 | - | - | 4 | 4 | 5 |
| **High and Dense Forests of the Orobiomes of Tropical Rainforest Zonobiome** | - | - | 0 | 3 | 5 | 0 | 4 | - | - | 4 | 4 | 5 |
| **High and Dense Forests of the Zonobiome of Tropical Sub-Humid Forests** | - | - | 0 | 3 | 5 | 0 | 4 | - | - | 4 | 4 | 5 |
| **High and Dense Forests of the Zonobiome of Tropical Rainforests** | - | - | 0 | 3 | 5 | 0 | 4 | - | - | 4 | 4 | 5 |
| **High and Dense Forests of the Zonobiome of Tropical Dry Forests** | - | - | 0 | 3 | 5 | 0 | 4 | - | - | 4 | 4 | 5 |
| **High and Dense Forests and Pantanos of the Helobiomes of the Tropical Rainforest zonobiome** | - | - | 0 | 3 | 5 | 0 | 4 | - | - | 4 | 4 | 5 |
| **High and Half Dense Forests of the Helobiomes of the Tropical Rainforest Zonobiome** | - | - | 0 | 3 | 5 | 0 | 4 | - | - | 4 | 4 | 5 |
| **Low sclerophyllous dense forests (high Amazonian Caatingas) of the Pedobiomes of the Zonobiome of the Tropical Rainforests** | - | - | 0 | 3 | 5 | 0 | 4 | - | - | 4 | 4 | 5 |
| **Dense low forests and mangrove shrublands of the Zonobiome Helobiomes of Tropical Rainforest** | - | - | 0 | 3 | 5 | 0 | 4 | - | - | 4 | 4 | 5 |
| **Dense low forests and Shrublands of the Zonal Orobiomes of the Zonobiome of Tropical Rain forest** | - | - | 0 | 3 | 5 | 0 | 4 | - | - | 4 | 4 | 5 |
| **Low Forests and Dense Shrublands of the Helobiomes of Zonobiomes of Tropical Rainforest** | - | - | 0 | 3 | 5 | 0 | 4 | - | - | 4 | 4 | 5 |
| **Low forests and dense shrublands of the Zonobiome of the Tropical Dry Forests** | - | - | 0 | 3 | 5 | 0 | 4 | - | - | 4 | 4 | 5 |
| **Low forests and dense sclerophyllous shrublands (Middle Amazonian Caatingas) of the Pedobiomes of the Zonobiome of the Tropical Rainforests** | - | - | 0 | 3 | 5 | 0 | 4 | - | - | 4 | 4 | 5 |
| **Low forests, grasslands and floating vegetation of the Helobiomes of the Zonobiome of Tropical Dry Forest** | - | - | 0 | 3 | 5 | 0 | 4 | - | - | 4 | 4 | 5 |
| **Potential Ecosystem** |  |  |  |  |  |  |  |  |  |  |  |  |
| **Medium Dense Forests of the Helobiomes of the Tropical Rainforest Zonobiome** | - | - | 0 | 3 | 5 | 0 | 4 | - | - | 4 | 4 | 5 |
| **Medium Dense Forests of the Orobiomes of the Tropical Rainforest Zonobiome** | - | - | 0 | 3 | 5 | 0 | 4 | - | - | 4 | 4 | 5 |
| **Medium Dense Forests of the Zonobiome of the Tropical Rainforest** | - | - | 0 | 3 | 5 | 0 | 4 | - | - | 4 | 4 | 5 |
| **Medium Dense Forests of the Zonobiome of the Tropical Dry Forests** | - | - | 0 | 3 | 5 | 0 | 4 | - | - | 4 | 4 | 5 |
| **Medium and Low Dense Forests of the Helobiome of the Zonobiome of Tropical Deserts** | - | - | 0 | 3 | 5 | 0 | 4 | - | - | 4 | 4 | 5 |
| **Medium and Low Dense Forests of the Helobiomes of the Tropical Rainforest Zonobiome** | - | - | 0 | 3 | 5 | 0 | 4 | - | - | 4 | 4 | 5 |
| **Forests, grasslands and swamps of Helobiomes O_ZBH** | - | - | 0 | 3 | 5 | 0 | 4 | - | - | 4 | 4 | 5 |
| **Grasslands (campinas amazonicas) of the Pedobiomes of the Zonobiome of the Tropical Rainforests** | - | - | - | - | 5 | 0 | 3 | 0 | 2 | 3 | 4 | 5 |
| **Open grasslands and snows of the Orobiomes of the Zonobiome of Tropical Rainforest** | - | - | - | - | 5 | 0 | 3 | 0 | 2 | 3 | 4 | 5 |
| **Grasslands of the Orobiomes of the Tropical rainforest Zonobiome** | - | - | 0 | 3 | 5 | 0 | 4 | 0 | 3 | 4 | 4 | 5 |
| **Grasslands and dense shrublands of the Orobiomes of the Zonobiome of Tropical Rainforest** | 0 | 4 | 0 | 3 | 5 | 0 | 4 | 0 | 3 | 4 | 4 | 5 |
| **Lakes, lagoons, permanent swamps, and major rivers of the Hydrobiomes** | - | - | - | - | 5 | 0 | 3 | - | - | 3 | 3 | 5 |
| **Savannas casmófitas and grasslands of the Pedobiomes of the Zonobiome of Tropical Rainforest** | - | - | - | - | 5 | 0 | 2 | 0 | 2 | 3 | 3 | 5 |
| **Herbaceous savannas with shrublands of the zonal orobiomes of the Zonobiome of the Tropical Rainforest** | 0 | 4 | 0 | 3 | 5 | 0 | 4 | 0 | 3 | 4 | 4 | 5 |
| **Herbaceous savannas with shrublands of the Pedo / Peinobiomes of the Dry Tropical Forest Zonobiome and BsHT** | 0 | 3 | - | - | 5 | 0 | 2 | 0 | 2 | 3 | 3 | 5 |
| **Herbaceous savannas and shrublands of the Pedobiomes of the Zonobiome of the Tropical Rainforests** | 0 | 3 | - | - | 5 | 0 | 2 | 0 | 2 | 3 | 3 | 5 |

**Results**

LHIF increment rates at the national level are presented in table A-3. These values were calculated as the percentage LFHI difference between periods. We find that 1970-1990 is the time period with the highest increase, corresponding to the period with the highest growth rate, while 2000-2015 is the period that presents lower values of increase. The increase of the footprint at national level corresponds to approximately 1% during the 45 years studied.

**Table A-3. Increase in L**HFI **values and annual growth rates of the footprint at the national level.**

| **Average L**HFI **value** | | | | **Percentage increase in LHFI** | | | | **Annual growth rate of LHFI** | | | |
| --- | --- | --- | --- | --- | --- | --- | --- | --- | --- | --- | --- |
| **1970** | **1990** | **2000** | **2015** | **% of change 1970-1990** | **% of change 1990-2000** | **% of change 2000-2015** | **% of change 1970-2015** | **% annual growth 1970-1990** | **% annual growth 1990-2000** | **% annual growth 2000-2015** | **% annual growth 1970-2015** |
| 15.36 | 19.71 | 21.39 | 23.04 | 28.32 | 8.52 | 7.71 | 50.00 | 1.25 | 0.82 | 0.50 | 0.91 |

Table A-4 shows the percentage increment of the mean LHFI for each natural region evaluated. We found that the highest increases occurred from the 1970s to 1990s, when the Orinoco region showed the greatest increase with 56%, followed by the Pacific region with 55%. When observing the total increase (1970-2015) we found that the Pacific region showed the highest increases with 148% followed by the Amazon region with 116%, the Caribbean region showed the lowest values of increase in all the periods evaluated.

| **Natural Region** | **Average** LHFI **value** | | | | **Percentage increase in LHFI** | | | |
| --- | --- | --- | --- | --- | --- | --- | --- | --- |
|  | **1970** | **1990** | **2000** | **2015** | **% Change 1970-1990** | **% Change 1990-2000** | **% Change 2000-2015** | **% Change 1970-2015** |
| Andean | 27.33 | 34.08 | 36.83 | 38.46 | 24.73 | 8.04 | 4.43 | 40.74 |
| Amazon | 3.17 | 4.60 | 5.44 | 6.85 | 45.31 | 18.20 | 25.94 | 116.29 |
| Orinoco | 10.37 | 16.18 | 17.95 | 20.14 | 56.00 | 10.91 | 12.24 | 94.22 |
| Caribbean | 39.98 | 46.55 | 48.48 | 49.84 | 16.43 | 4.13 | 2.81 | 24.65 |
| Pacific | 3.96 | 6.14 | 7.15 | 9.83 | 55.01 | 16.38 | 37.61 | 148.24 |
| Catatumbo | 26.12 | 30.12 | 33.95 | 35.84 | 15.31 | 12.71 | 5.57 | 37.20 |

**Table A-4.** Increase in LHFI values for each region during the evaluated periods

Table A-5 shows the annual growth rate of the human footprint for each region evaluated. For all the periods evaluated, the growth rates were maintained between 0.1 and 3%; the Amazon, Orinoco and Pacific regions showed the highest growth rates in all periods, while the Andean, Catatumbo and Caribbean regions had the lowest values.

| **Natural Region** | **Annual growth rate of LHFI** | | | |
| --- | --- | --- | --- | --- |
|  | **% Annual growth**  **1970-1990** | **% Annual growth**  **1990-2000** | **% Annual growth**  **2000-2015** | **% Annual growth**  **1970-2015** |
| Andean | 1.11 | 0.78 | 0.29 | 0.76 |
| Amazon | 1.89 | 1.69 | 1.55 | 1.73 |
| Orinoco | 2.25 | 1.04 | 0.77 | 1.49 |
| Caribbean | 0.76 | 0.41 | 0.19 | 0.49 |
| Pacific | 2.22 | 1.53 | 2.15 | 2.04 |
| Catatumbo | 0.71 | 1.20 | 0.36 | 0.71 |

**Table A-5.** Rates of annual growth of LHFI at the regional level.

IDEAM, 2010. Land Cover Map Corine Land Cover Methodology Adapted for Colombia. Scale1:100.000. Instituto de Hidrología, Meteorología y Estudios Ambientales. Bogotá, D. C., 72p.

IGAC, 2016. Base De Datos Cartográfica Escala, 1:100 000, proyecto: Carta general a escala 1:100000, VERSION 2014_1. 2016.

Tapia-Armijos, M.F., Homeier, J., Draper Munt, D., 2017. Spatio-temporal analysis of the human footprint in South Ecuador: Influence of human pressure on ecosystems and effectiveness of protected areas. Appl. Geogr. 78, 22–32. https://doi.org/10.1016/j.apgeog.2016.10.007

The University of Texas at Austin., 2015. Perry-Castañeda Library, Map Collection-Colombia Maps” Disponible en: <https://legacy.lib.utexas.edu/maps/colombia.html>. Ultimo acceso: 22 de marzo de 2019.

Woolmer, G., Trombulak, S.C., Ray, J.C., Doran, P.J., Anderson, M.G., Baldwin, R.F.,Morgan, A., Sanderson, E.W., 2008. Rescaling the human footprint: a tool forconservation planning at an ecoregional scale. Landscape Urban Plann. 87 (1),42–53, http://dx.doi.org/10.1016/j.landurbplan.2008.04.005.
